# PIWI silencing mechanism involving the retrotransposon *nimbus* orchestrates resistance to infection with *Schistosoma mansoni* in the snail vector, *Biomphalaria glabrata*

**DOI:** 10.1101/2021.01.12.426235

**Authors:** Michael Smith, Swara Yadav, Olayemi Akinyele, Nana Adjoa Pels, Daniel Horton, Nashwah Alsultan, Andrea Borns, Carolyn Cousin, Freddie Dixon, Victoria H. Mann, Clarence Lee, Paul J. Brindley, Najib M. El-Sayed, Joanna M. Bridger, Matty Knight

## Abstract

**Background:** Schistosomiasis remains widespread in many regions despite efforts at its elimination. By examining changes in the transcriptome at the host-pathogen interface in the snail *Biomphalaria glabrata* and the blood fluke *Schistosoma mansoni,* we previously demonstrated that an early stress response in juvenile snails, manifested by induction of heat shock protein 70 (Hsp 70) and Hsp 90 and of the reverse transcriptase (RT) domain of the *B. glabrata* non-LTR-retrotransposon, *nimbus*, were critical for *B. glabrata* susceptibility to *S. mansoni*. Subsequently, juvenile *B. glabrata* BS90 snails, resistant to *S. mansoni* at 25°C become susceptible by the F2 generation when maintained at 32°C, indicating an epigenetic response.

**Methodology/Principal Findings:** To better understand this plasticity in susceptibility of the BS90 snail, mRNA sequences were examined from *S. mansoni* exposed juvenile BS90 snails cultured either at 25°C (permissive temperature) or 32°C (non-permissive). Comparative analysis of transcriptomes from snails cultured at the non-permissive and permissive temperatures revealed that whereas stress related transcripts dominated the transcriptome of susceptible BS90 juvenile snails at 32°C, transcripts encoding proteins with a role in epigenetics, such as PIWI (*BgPiwi*), chromobox protein homolog 1 (*BgCBx1*), histone acetyl transferase histone deacetylase (HDAC) and metallotransferase (MT) were highly expressed in those cultured at 25°C. To further determine a role for *BgPiwi* in *B. glabrata* susceptibility to *S. mansoni*, siRNA corresponding to the *BgPiwi* encoding transcript was utilized to suppress expression of *BgPiwi*, rendering the resistant BS90 juvenile snail susceptible to infection at 25°C. Given transposon silencing activity of PIWI as a facet of its role as guardian of the integrity of the genome, we examined the expression of the *nimbus* RT encoding transcript at 120 min after infection of resistant BS90 *piwi-*siRNA treated snails. We observed that *nimbus* RT was upregulated, indicating that modulation of the transcription of the *nimbus* RT was associated with susceptibility to *S. mansoni* in *BgPiwi-* siRNA treated BS90 snails. Furthermore, treatment of susceptible snails with the RT inhibitor lamivudine, before exposure to *S. mansoni*, blocked *S. mansoni* infection concurrent with downregulation of the *nimbus* RT transcript and upregulation of the *BgPiwi* encoding transcript in the lamivudine-treated, schistosome-exposed susceptible snails.

**Conclusions and Significance:** These findings support a role for the interplay of *BgPiwi* and *nimbus* in the epigenetic modulation of plasticity of resistance/susceptibility in the snail-schistosome relationship.

## INTRODUCTION

The freshwater snail, *Biomphalaria glabrata,* is an obligate intermediate host of the trematode, *Schistosoma mansoni*, the causative agent of the neglected tropical disease (NTD) schistosomiasis in neotropical regions. At least 600 million people, mainly in subSaharan Africa, are at risk for schistosomiasis, a number that remains excessively high, despite efforts to control transmission of the disease [1]. This disease causes widespread chronic morbidity and male and female infertility. Specifically, infections caused by the species, *Schistosoma haematobium* may result in bladder cancer and female genital schistosomiasis. The latter exacerbates transmission of sexually transmitted diseases including HIV (AIDS) [2].

The disease burden from schistosomiasis is probably underestimated, and it has been suggested that the number of infected individuals exceeds 400 million [3]. On the other hand, there are other estimations that claim 230 million people worldwide are infected with *S. mansoni* [1]. These numbers are often underestimated due to the inability of current diagnostic methods to detect light infections [4]. There are few defenses against schistosomiasis, mainly because the residents of infected areas lack sufficient infrastructure to properly combat this disease [3]. An integrated control approach, implementing mass chemotherapy and molluscicides has made a difference in breaking the complex life cycle of the parasite but without new effective drugs and vaccines to prevent re-infection in treated areas, the cycle of repeated infection is the norm, thereby making the long-term control of schistosomiasis elusive [5, 6]. Because of recent projections made by the World Health Organization to eliminate schistosomiasis by 2025 and coupled with recent concerns of the spread of the disease into Mediterranean countries of Europe, alternative approaches focusing mainly on blocking the development of the parasite in the snail host are aggressively sought [7–9].

Molecular mechanism(s) involved in shaping the relationship between the parasite and its obligate intermediate snail host, *Biomphalaria glabrata*, remains largely unknown. However, information that will assist in clarifying some of the mechanisms of host/parasite interactions is steadily being amassed. Reference genome sequences for all three organisms (human, schistosome and snail) pertinent to transmission of schistosomiasis are now available [10–12]. Additionally, genes that underlie resistance and susceptibility phenotypes in *B. glabrata* to *S. mansoni* infection are being identified [13–15], as are transcripts encoding larval (miracidia) parasite proteins that are expressed at the snail/parasite interphase [16, 17]. Both snail and parasite determinants are involved in a complex and dynamic innate defence system that either rejects or sustains the successful development of the intra-molluscan stages of the parasite [13, 15].

Previously, we demonstrated that juvenile *B. glabrata*, that are either resistant or susceptible to *S. mansoni,* display a differential stress response after early exposure to wild type but not to irradiated *S. mansoni* miracidia. The stress response observed in the susceptible juvenile snail was manifested by the early induction of transcripts encoding heat shock proteins (Hsp) 70, Hsp90 and the reverse transcriptase (RT) domain of the *B. glabrata* non-LTR-retrotransposon, *nimbus* [18]. Furthermore, the non-random relocalization of the Hsp70 gene loci in interphase nuclei preceded transcription of the corresponding Hsp70 transcript in the susceptible but not in the resistant snail, indicating that in-coming schistosomes possess the ability to orchestrate in a rapid and systemic fashion, the genome remodeling of juvenile susceptible snails soon after infection [19]. We also demonstrated that resistance in the juvenile BS90 snail stock was a temperature dependent trait. Thus, when cultured at room temperature (25°C), juvenile BS90 snails remained consistently resistant to infection. However, when cultured at 32°C for several generations (F_I_ to F_3_), the progeny juvenile snails were phenotypically susceptible [20], indicating a plastic epigenetic control over resistance.

Other studies that have used adult (>7 mm in diameter) snails instead of juveniles have suggested that ability to alter the resistance of the BS90 to infection at elevated temperature might be a strain-specific trait [21]. In recent studies, resistant BS90 snails were found to be susceptible when exposed to *S. mansoni* as neonates [22]. In general, adult snails are less susceptible to infection [23]. Furthermore, in some stocks of *B. glabrata*, for example the 93375 strain, the juveniles are susceptible but become resistant to the same strain of *S. mansoni* as young adults (at the onset of fecundity), and once egg laying ceases and amoebocyte accumulations disappear in the pericardial wall, revert to the susceptible phenotype [23, 24].

To further investigate the molecular basis of susceptibility plasticity, notably in the BS90 snail, we have undertaken a comparative analysis of the transcriptomes of juvenile BS90 snails cultured for several generations at either permissive (32°C) or non- permissive (25°C), aiming to obtain leads for pathway(s) that lead either to resistance or susceptibility. This investigation and the findings are detailed below.

## MATERIALS & METHODS

### Snails

The BS-90 snail is a wild-type pigmented snail that is resistant to *S. mansoni* (NMRI strain) either as a juvenile or as an adult snail at 25°C [25]. The NMRI snail is an albino susceptible snail that was derived from a cross between the wild type Puerto Rican snail and a highly susceptible Brazilian albino snail [26, 27]. The BB02 snail is a susceptible pigmented wild type snail from Brazil whose genomic DNA sequence was recently reported [10]. The susceptible snails (NMRI and BBO2) are highly susceptible as juveniles but the degree of susceptibility, especially in the NMRI stock, is variable as an adult snail [28].

### Snail husbandry and *S. mansoni* infections

BS90 stocks were cultured at 32°C as described [20]. Exposure of BS90 snails cultured at 32°C to miracidia were performed using juvenile progeny, F_1_-F_2_, <4 mm in diameter that were bred at the elevated temperature. Briefly, BS90 snails were cultured either at 25°C or 32°C in freshly made artificial pond water (www. afbr-bri.com) and fed *ad libitum* with either romaine lettuce or snail gel food [29]. Juvenile snails or egg clutches were transferred from 25°C to 32°C and maintained in groups of 3 or 4 in fresh water (100 ml) in beakers maintained in a water bath at 32°C. The temperature inside the water bath was monitored daily to maintain 32°C for the duration of the experiment. The snails were cleaned weekly making sure that pre-warmed (32°C) water was used to clean the snails. Detritus including dead snails and decayed lettuce leaves were removed daily. The egg clutches from these snails (produced at 32°C) were collected and their progeny were maintained at 32°C until they had grown to 3 to 4 mm in diameter (juvenile snails) before exposure to miracidia at 25°C. The juvenile BS90 snails, 3 to 4 mm in diameter, were maintained for two generations at either 32°C or 25°C and RNA prepared from 0 and two-hour infected F2 progeny as described [20]. Snails were exposed individually to the 10 to 12 miracidia in wells of a 12-well tissue culture plate (Greiner Bio-One, North Carolina, USA) at room temperature. Miracidia were hatched from eggs recovered from the livers of mice which had been infected with *S. mansoni* (NMRI strain) for seven weeks [30].

Exposed snails (not used for RNA) were maintained at 25°C and examined for cercarial shedding from four to 10 weeks later. Susceptible BBO2 and NMRI *B. glabrata* snails utilized in this study were exposed as juveniles (described above) to freshly hatched miracidia. To determine patency (cercarial shedding), individual snails were immersed in one ml nuclease-free water in 12-well plates and directly exposed to a light source for 30 to 60 minutes at room temperature, after which snails were removed from the wells. Cercariae released from individual snails were counted after adding a few drops of Lugol’s iodine solution to each well. After shedding, snails that were patent were euthanized by immersion in 95% ethanol; non-shedding snails were incubated for up to 10 weeks at 25°C and checked weekly for patency.

### RNA sequencing, assembly, and annotation

Total RNA was isolated by RNAzol (Molecular Research Center, Inc. Cincinnati, OH) from resistant (25°C) and susceptible (32°C) BS90 snails. BS90 snail transcriptome was generated from polyA^+^ RNA isolated from pooled intact 2 hour exposed juvenile snails maintained either at the non-permissive temperature (25°C), or at the permissive temperature (32°C) on an Illumina HiSeq 100. Following RNAseq Illumina sequencing, *de novo* assembly of the transcriptome was performed using Trinity. Functional annotation of the contigs was performed with Trinotate that included Gene Ontology assignments when possible, pFAM domain identification, transmembrane region predictions (TmHMM) and signal peptide predictions (signalP). Differential expression (DE) analyses were performed by using DESeq and identified differentially expressed contigs between BS90 snails from the different categories. The analyses of these contigs revealed the presence of known specific genes coding for several stress related and other transcripts (Table 1). The complete transcriptome profiles of changes observed in schistosome juvenile BS90 snails at either permissive (32°C) or non-permissive (25°C) are provided as FASTQ files in the SRA database within NCBI with Bioproject ID PRJNA687288.

### Two step qPCR

Differential expression of the selected transcripts identified from the single pass RNAseq dataset generated from resistant (25°C) and susceptible (32°C) juvenile BS90 snails were further validated in other representative BS90 and susceptible snail stocks NMRI and BBO2 that were either unexposed (0 hour) or exposed to *S. mansoni* for 30 min, 1, 2, 4 and 16 hours at 25°C. cDNA was prepared from total RNA as described and contaminating genomic DNA removed by treatment of the RNAs with DNase (RQI Promega WI) before performing the RT qPCR assays (Ittiprasert, 2012 #7020). Quantitative real time PCR was performed with forward and reverse gene specific primers corresponding to selected transcripts normalized against the expression of the myoglobin gene as a reference, as described [31]. Oligonucleotide primers (forward and reverse, Table 1) for transcripts identified by sequencing RNA of BS90 snails at either permissive (32°C) or non-permissive temperatures (25°C) were designed from amino acid sequences corresponding to the coding DNA sequence (CDS) of the following transcripts: piwi like protein (*BgPiwi*), chromobox protein homolog 1 (*BgCBx1*), methyltransferase (*BgMT*), Histone Acetyl Transferase (*BgHAT*) and histone deacetylase (*BgHDAC*). These CDS were utilized to interrogate the reference *B. glabrata* genome sequences in GenBank by using the Basic Local Alignment Search Tool, BLAST in NCBI. A standard BLASTp was performed to further validate annotations of the selected transcripts followed by a SMART BLAST to explore the phylogeny of the *B. glabrata* orthologs to other transcripts in the public domain. Amino acid sequences of *B. glabrata* CDS showing significant (E value = < 10^-4^ and with > 25% amino acid sequence identity) homology to other transcripts were converted to the nucleotide sequences and gene specific primers were designed using primer BLAST. Forward and reverse primers for qPCR analysis were obtained from Eurofins Genomics (Louisville, KY) after the exclusion of sequences for *S. mansoni* to avoid any possible amplification of parasite RNA during Real Time PCR analysis. Two-step RT qPCR was utilized to quantitatively assess expression of the selected transcripts using 500 ng of cDNA as template. SYBR Green PCR Master Mix kit (Applied Bio systems, Thermo Fisher Scientific, Wolston Warrington, UK), with 15 μM of forward and reverse primers were used to evaluate the temporal expression of PIWI, Chromobox protein homolog 1, HDAC, HAT and methyl transferase (MT). Each sample was run in triplicate and reactions normalized against the constitutively expressed myoglobin reference gene in a 7300-thermal cycler (Applied Biosystems). Relative quantitative expression of the genes of interest between resistant and susceptible snails was evaluated by the ΔΔCt method. The resulting fold change in expression of the genes of interest normalized against the signal for myoglobin were calculated by using the formula, 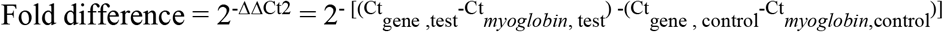 [32]. Differences were assessed using Student’s *t* test, Welch’s *t* test and 2-way analysis of variance (ANOVA) wherever relevant by comparing the differential expression (delta-Ct value) of the transcripts among treatment and control groups. A *p*-value of <0.05 was considered to be statistically significant, with level of significance denoted as follows, ****, *p* ≤ 0.0001, ***, *p* ≤ 0.001, **, *p* ≤ 0.01, *, *p* ≤ 0.05, and ns, *p*> 0.05

### *BgPIWI* transcript silencing by dsRNA and siRNA

To investigate the functional role of *Bgpiwi* expression in *B. glabrata* susceptibility to schistosome infection, the transcription of *Bgpiwi* was silenced by soaking juvenile BS90 snails in either dsRNA-, or siRNA-PEI complexes [33]. Double–stranded (ds)RNA corresponding to *Bgpiwi* was synthesized by using an *in vitro* transcription kit with a purified *Bgpiwi* PCR product containing T7 sequences (sense and antisense) as template according to the manufacturer’s instructions (MEGAscriptT7, ThermoFisher Scientific Inc.) [34]. Off-target silencing of the transcript encoding *Bgpiwi* in the resistant BS90 snail was evaluated by soaking snails in parallel in universal mock siRNA-PEI complexes as control (MISSION siRNA Universal negative control#1, Sigma Aldrich, St. Louis, MO). Knock-down of the *Bgpiwi* transcript in the resistant BS90 snail (cultured at 25°C) was done as follows: juvenile snails were placed in 1.0 ml nuclease-free dH_2_O containing either 300 ng dsRNA: 1.0 μg PEI nanoparticle complexes or 775 ng siRNA: 1.0 μg PEI nanoparticle complexes. The complexes were prepared as follows: in a 1.5 ml capacity microcentrifuge tube, 1 μg of PEI, branched with average molecular weight 25000, (Sigma Aldrich) in 500 μl nuclease-free H_2_O was added slowly, drop-wise, to two different siRNAs (Sigma Millipore) (start on target sequence CDS position 2393bp-sense: GAACCAUUGUGGAUCAAAU/anti-sense: AUUUGAUCCACAAUGGUUC; (start on target sequence at CDS position 2403bp sense: GGAUCAAAUAAUUACGAA/anti-sense: UUUCGUAAUUAUUUGAUCC) diluted in 500 μl before mixing vigorously for 10 seconds at room temperature. Both duplex siRNAs corresponding to *BgPiwi* transcript were utilized simultaneously in a single tube. Samples of siRNA/PEI complexes (Total of 1.0 ml in microcentrifuge tubes) were incubated at room temperature for 20 minutes before placing individual juvenile snails in the mixtures. Holes were punched in lids of the closed microcentrifuge tubes containing snails in siRNA/PEI complexes before incubating overnight at room temperature. Control tubes, incubated in parallel, contained the following samples, a) *Bgpiwi* siRNAs without PEI, b) PEI only without *Bgpiwi* siRNAs and c) Mission Universal mock siRNA/PEI complexes (Sigma Millipore). RNA was isolated as described above from washed transfected snails before utilizing for qPCR as described above. For each assay, quantitative expressions of *Bgpiwi* and *nimbus*RT transcripts (normalized against myoglobin expression) were evaluated with and without *S. mansoni* infection by qPCR using forward and reverse primers corresponding to either transcript (Table 1). Transfected snails (with and without infection) that were not investigated by RNA-based assays were maintained in at 25°C as above and monitored for cercarial shedding at 4, 6, or 10 weeks later. The silencing of *Bgpiwi* with dsRNA/PEI was evaluated in three, and with siRNA/PEI, in four biological replicates, respectively.

### Treatment of susceptible BBO2 snails with reverse transcriptase inhibitor, lamivudine

Given that the central role of PIWI involves silencing of endogenous mobile elements, such as nimbus, a non-LTR retrotransposon in the genome of *B. glabrata* [10, 35], we examined the modulation of expression of the transcript encoding the RT domain of *nimbus* in the following categories of susceptible BBO2 snails – a) normal snails, b) snails treated overnight at room temperature with lamivudine at 100 ng/ml (Sigma Aldrich, St. Louis, MO) and c) BS90 resistant snails treated with siRNA corresponding to *Bgpiwi/*PEI complexes, as described above. Snails in these categories (a to c) were either unexposed (0) or exposed (individually) to 10 miracidia for 2 hours at 25°C. Before exposure, individual snails incubated in either lamivudine or *Bgpiwi* siRNA/PEI complexes were washed twice with water before transfer to 2.0 ml water in 6-well tissue culture plates to which freshly hatched miracidia (isolated from 7 weeks infected mouse liver homogenate) was added and maintained for 120 min at room temperature. Exposed and unexposed snails from either lamivudine-treated susceptible BBO2 or BS90 *siRNABgPiwi/*PEI-treated snails were either frozen immediately at −80°C in RNAzol until required for RNA isolation or, if not used for RNA preparation immediately, transferred into 500 ml beakers containing aerated tap water and maintained as described above at room temperature and evaluated for cercarial shedding at week 4, 6 or 10 postexposure. For comparison, susceptible snails were also either pre-treated as described above, or after two weeks post-exposure, with another RT inhibitor BPPA (Santa Cruz Biotechnology Inc., CA) that specifically inhibits the catalytic RT domain of human telomerase (hTERT).

### Examining genome organization, relocation of the *piwi* locus, in susceptible and resistant snails following exposure to *S. mansoni* miracidia

Fluorescence *in situ* hybridisation (FISH) was performed using a probe derived from *B. glabrata* bacterial artificial chromosome (BAC) libraries for the *piwi* locus. The DNA probe was labelled by nick translation (BioNick Invitrogen, UK) as described [36, 37] and incubating for 45-50 mins. The probe was precipitated with 1μg of labelled BAC DNA)[36], 80 μg of *B. glabrata* genomic DNA and 9 μg of herring sperm DNA. These components were dissolved in 48 μL of hybridisation mix at room temperature overnight, this amount can be used for up to four slides. Preparation and fixation of samples followed the protocol described previously [19]. Snail shells were crushed using a microscope slide and the ovotestes excised using needle-nose forceps. Each ovotestis was placed in a microcentrifuge tube containing 0.05 M KCl, macerated using a tissue grinder (Axygen, UK) and incubated in solution for 30 min at room temperature. Samples were then centrifuged at 200g for 5 min and supernatant discarded. Methanol:acetic acid [3:1, v/v] was presented dropwise, with agitation, to the samples. Once 0.5 mL of fixative was added, the samples were incubated at room temperature for 10 min before centrifuging again and discarding the supernatant, this fixation step was repeated twice with the final fix volume being 100 μL. Slides were also prepared by misting the slide with water vapour and then dropping 20 μL of a sample from a height onto the slide and allowing the slide to dry on a slide drier. The slides were aged by placing into a 70°C oven for 60 min then were taken through a dehydration series of 70%, 90% and 100% ethanol, spending 5 min in each solution. Slides were dried and warmed up to 37°C on a slide dryer alongside 22×22 coverslips in preparation for probe addition. Probe denaturation was performed at 75°C for 5 min and then allowed to reanneal for 20 min at 37°C before use. Hybridisation of samples and probe was performed using the Top Brite automatic slide hybridiser (Resnova, Italy). Eleven μL of probe was presented to a coverslip and the slide with the sample brought to the coverslip and the coverslip sealed to the slide using rubber cement (Weldtite). The Top Brite was set for 37°C for 2 min up to 75°C for 2 min and then lowered to 37°C for 30 min, once the slides had returned to 37°C they were transferred to a humidified chamber at 37°C for 72 hours. Post hybridisation, the rubber cement was removed and coverslips allowed to detach in the first wash. The washes were performed at 42°C in 2x SSC three times for 5 min each. A blocking solution, made of 4% BSA (Sigma Aldrich, UK) in 2x SSC, was prepared. After slides were removed from the third wash the excess solution was drained and 100 μL of blocking solution was added and the slides covered with parafilm. Slides were maintained in a humidified chamber at room temperature for 30 min. Streptavidin-Cy3 was diluted 1:200 in 1% BSA in 2x SSC. After the blocking solution was removed, 100 μL of streptavidin-Cy3 solution was placed on each slide, covered in parafilm and incubated in a humidified chamber at 37°C for 30 min. After the streptavidin-Cy3 incubation the slides were washed sequentially in 2x SSC for 5 min, 1x PBS + 0.1% Tween 20 (Sigma Aldrich) for 1 min, and 1x PBS for 1 min. Lastly, the slides were rinsed in sterile water before counterstaining with 4’,6-diamidino-2-phenylindole (DAPI) in mountant (H1200, Vectorshield).

### Image Analysis

Images of nuclei were captured either with an Olympus BX41 fluorescence microscope with a greyscale digital camera (Digital Scientific, UK) and the Smart Capture 3 software (Digital Scientific, UK) or a Leica DM4000 using a Leica DFC365 FC camera and the Leica Application Suite (LAS) imaging software. At least 50 nuclei were imaged for each condition and processed via erosion script analysis [38] [39] to assess gene loci positioning by using greyscale images and measuring the intensity of DAPI and fluorescence in situ hybridization (FISH) signal, using the DAPI to outline the nuclei to create five shells of equal area so that the intensity of the DAPI and FISH signal can be measured and the FISH signal normalised for position by dividing by the DAPI signal, averaged for the 50 images. Unpaired, equal variance, Student’s *t* tests were performed to ascertain significant differences in gene loci positioning within the different shells.

## RESULTS

### Transcripts encoding PIWI (*BgPiwi*), HDAC (*BgHDAC*), chromobox protein homolog 1(*BgCBx1*), histone acetyl transferase (*BgHAT*) and metallotransferase (*BgMT*) are differentially regulated in resistant BS90 and susceptible BBO2 snails

In order to confirm and validate results of differential expression (DE) obtained from the De-novo single-pass sequencing of RNA isolated from *S. mansoni* exposed juvenile BS90 F2 snails, which were cultured either at non-permissive (25°C), or permissive (32°C), temperatures. We investigated the expression of selected transcripts that have a role in epigenetics by two-step qRT-PCR performed with RNA isolated from several different individual juvenile *B. glabrata* snails that were either resistant (BS90 cultured at 25°C) or susceptible (BBO2 and BS90 cultured at 32°C) to *S. mansoni* infection. The investigation of temporal (0, 30 min, 1, 2, 4, and 16 hours) expression of *BgPiwi* in either juvenile resistant BS90 (25°C) or juvenile susceptible BBO2 following exposure to *S. mansoni* by qPCR showed that the transcript encoding the *B. glabrata* Piwi-like protein (Isoform 1 Accession number XP_013081375) was upregulated between 30 min and 2 hour (2-to 7-fold) post-exposure in the resistant BS90 but not in the susceptible BBO2 snail (Fig. 1A). Figure 1B shows the temporal expression of the transcript encoding *BgHDAC* (Accession Xp_0130754221.1). Results likewise showed upregulation of this transcript (1.8-fold) in the juvenile resistant BS90 but not in the juvenile susceptible (BBO2) snails 2 hours after *S. mansoni* exposure. Similarly, as shown in Figures 1C to 1E, results demonstrated that transcripts encoding, the chromobox protein homolog 1 (Fig. 1C, *BgCBx1*), histone acetyl transferase (Fig. 1D, *BgHAT*) and metallotransferase (Fig. 1E, *BgMT*) were also upregulated in BS90 resistant but not the susceptible BBO2 snails, following *S. mansoni* infection. Inductions of up to 13-and 10-fold for transcripts encoding the chromobox protein homolog 1 and MT, respectively (between 2 and 4 hours post exposure) were observed in exposed juvenile resistant BS90 but not their exposed juvenile BBO2 susceptible counterparts.

**Figure 1A.**
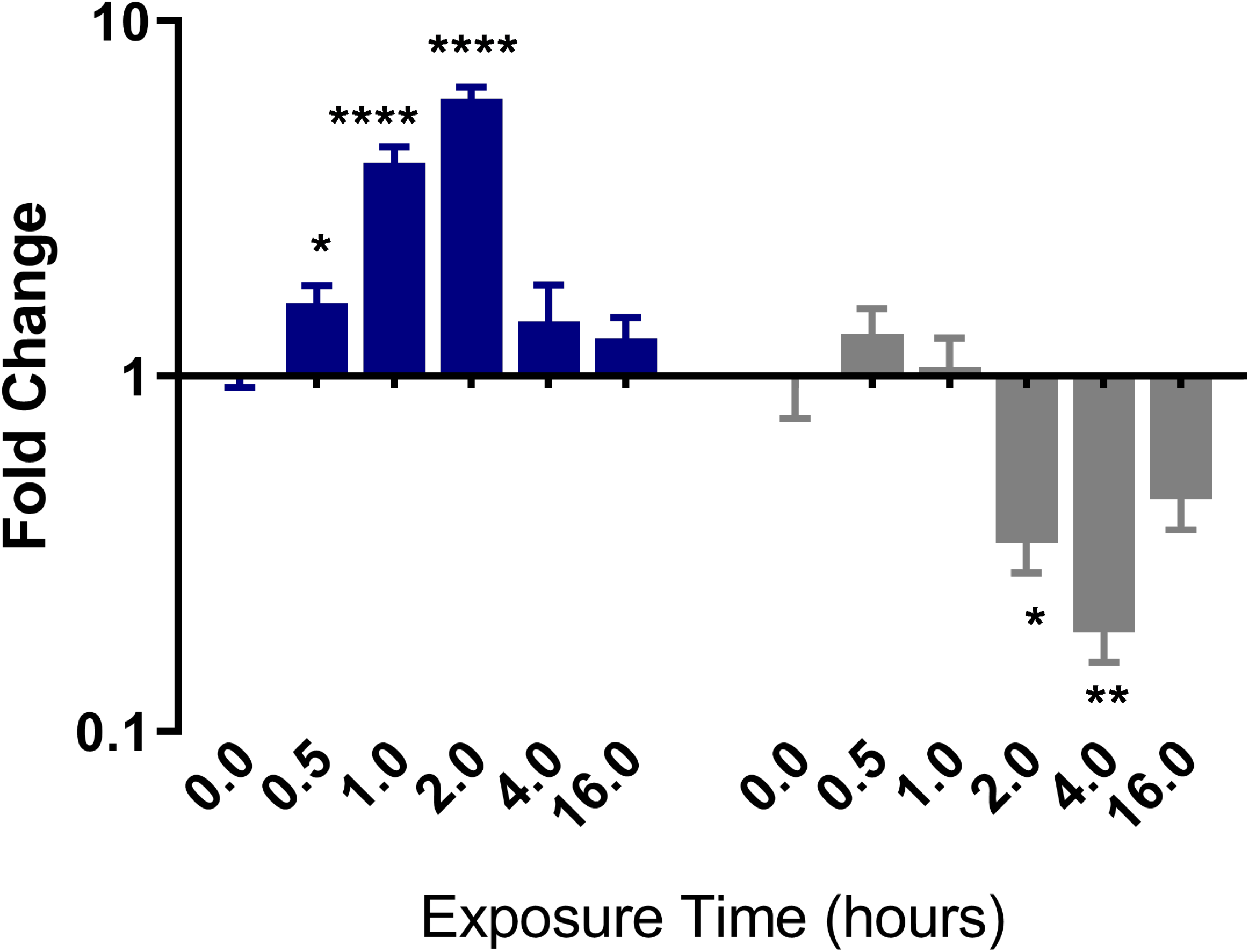
qPCR analysis of RNA from resistant BS-90 (blue histogram) or susceptible BBO2 (gray histograms) juvenile snails unexposed (0) or exposed for increasing intervals (30 seconds to 16 hours) to *S. mansoni* miracidia. Histograms show expression of the *BgPiwi* encoding transcript in snails at each time point from five biological replicates. Note the increase in fold change in the resistant BS-90 compared to the susceptible BBO2 snails after parasite infection. Significant expression normalized against expression of the myoglobin encoding transcript was measured by 2-way ANOVA and is indicated by number of asterixis on each histogram where ****, indicates the most significant value *p* ≤ 0.0001, *** *p* ≤ 0.001, ** *p* ≤ 0.01, * *p* ≤ 0.05, ns *p* > 0.05.

**Figure 1B.**
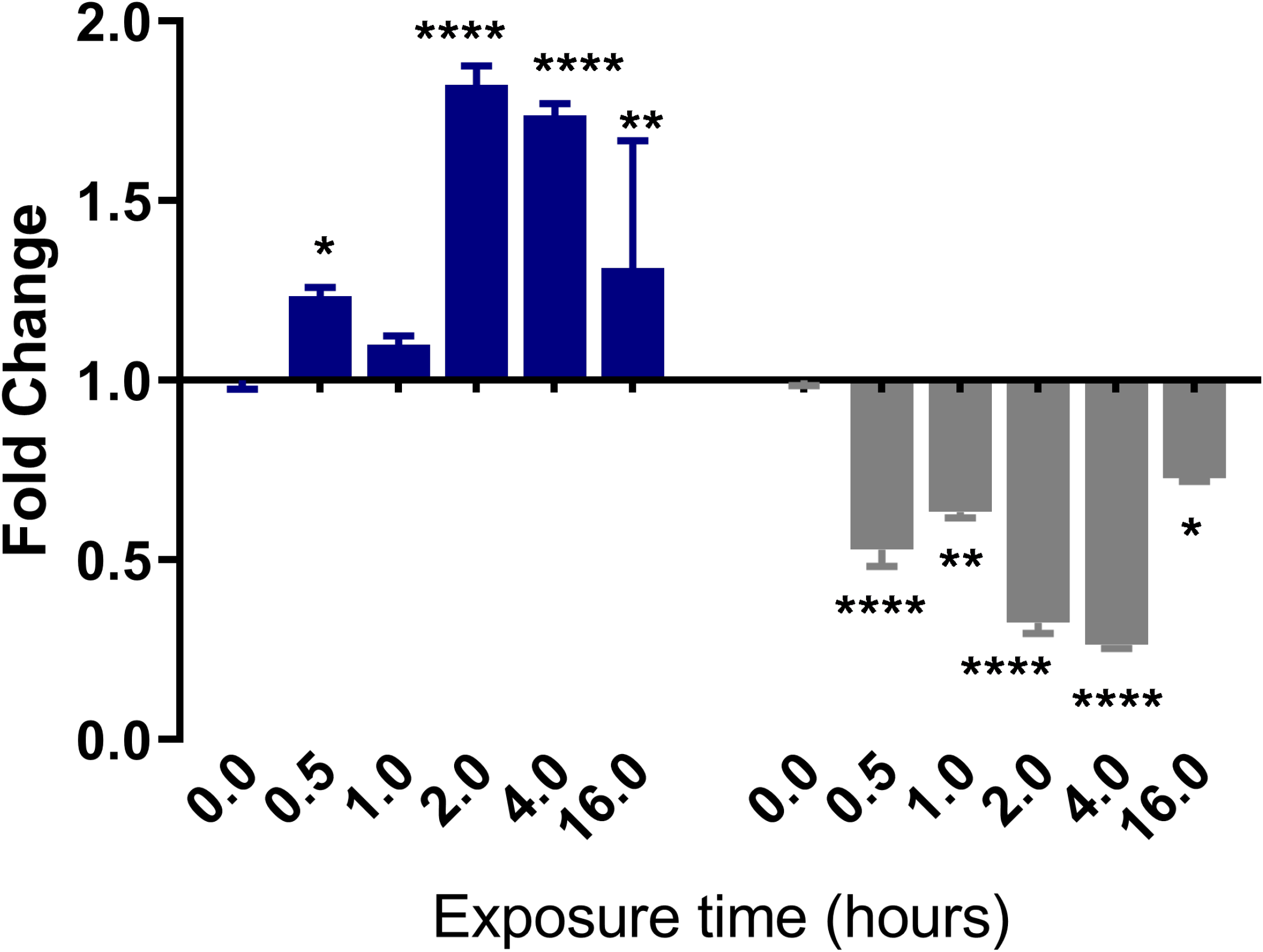
qPCR analysis of RNA from either resistant BS-90 (blue histogram) or susceptible BBO2 (gray histograms) juvenile snails unexposed (0) or exposed for increasing intervals (30 seconds to 16 hours) to *S. mansoni* miracidia. Histograms show expression of the *BgHDAC* encoding transcript in snails at each time point from five biological replicates. Note the increase in fold change in the resistant BS-90 compared to the susceptible BBO2 snails after parasite infection. Significant expression normalized against expression of the myoglobin encoding transcript was measured by 2-way ANOVA and is indicated by number of asterixis on each histogram where ****, indicates the most significant value *p* ≤ 0.0001, *** *p* ≤ 0.001, ** *p* ≤ 0.01, * *p* ≤ 0.05, ns *p* > 0.05.

**Figure 1C.**
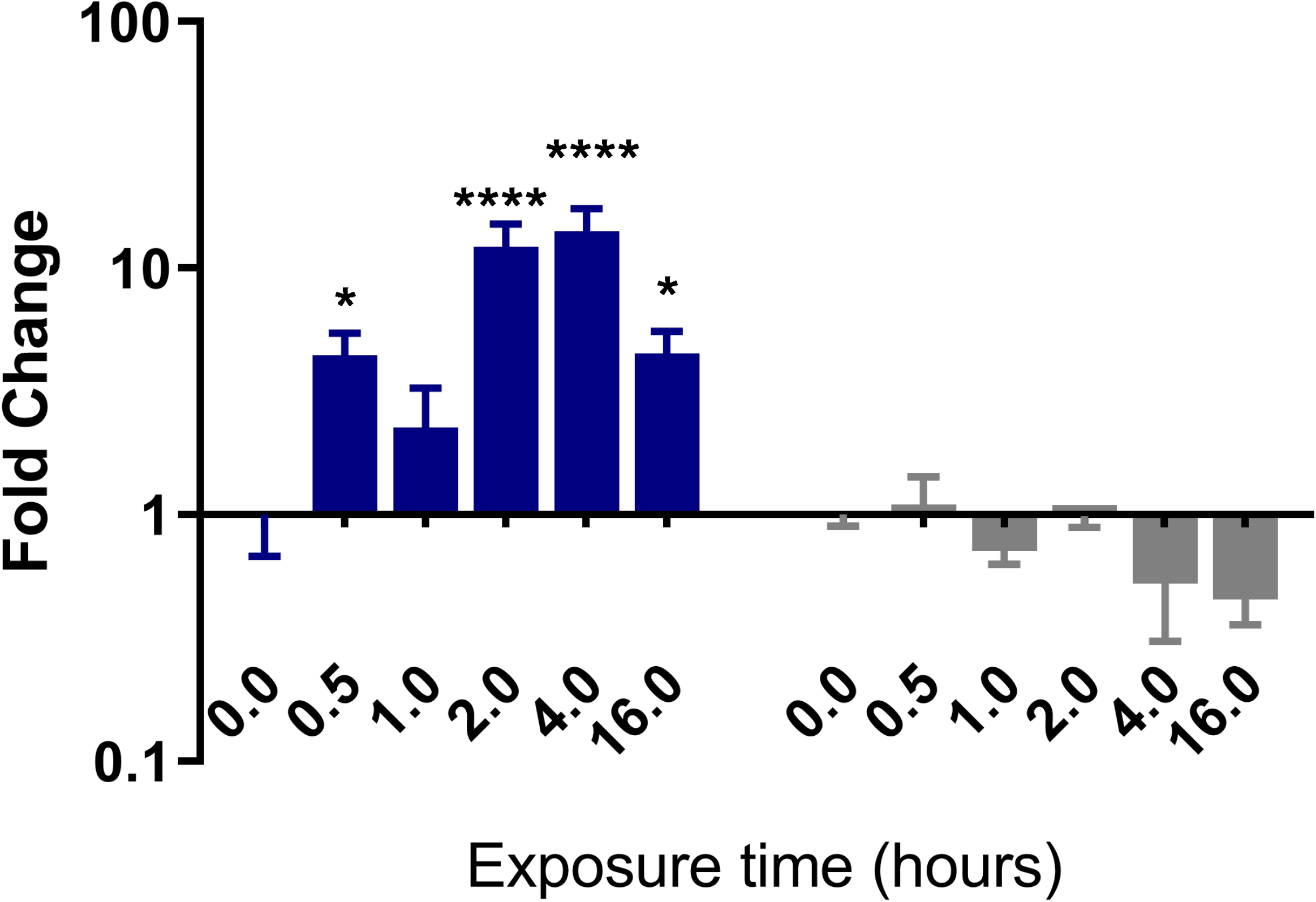
qPCR analysis of RNA from either resistant BS-90 (blue histogram) or susceptible BBO2 (gray histograms) juvenile snails unexposed (0) or exposed for increasing intervals (30 seconds to 16 hours) *S. mansoni* miracidia. Histograms show expression of the *BgCBx* encoding transcript in snails at each time point from five biological replicates. Note the increase in fold change in the resistant BS-90 compared to the susceptible BBO2 snails after parasite infection. Significant expression normalized against expression of the myoglobin encoding transcript was measured by 2-way ANOVA and is indicated by number of asterixis on each histogram where ****, indicates the most significant value *p* ≤ 0.0001, *** *p* ≤ 0.001, ** *p* ≤ 0.01, * *p* ≤ 0.05, ns *p* > 0.05.

**Figure 1D.**
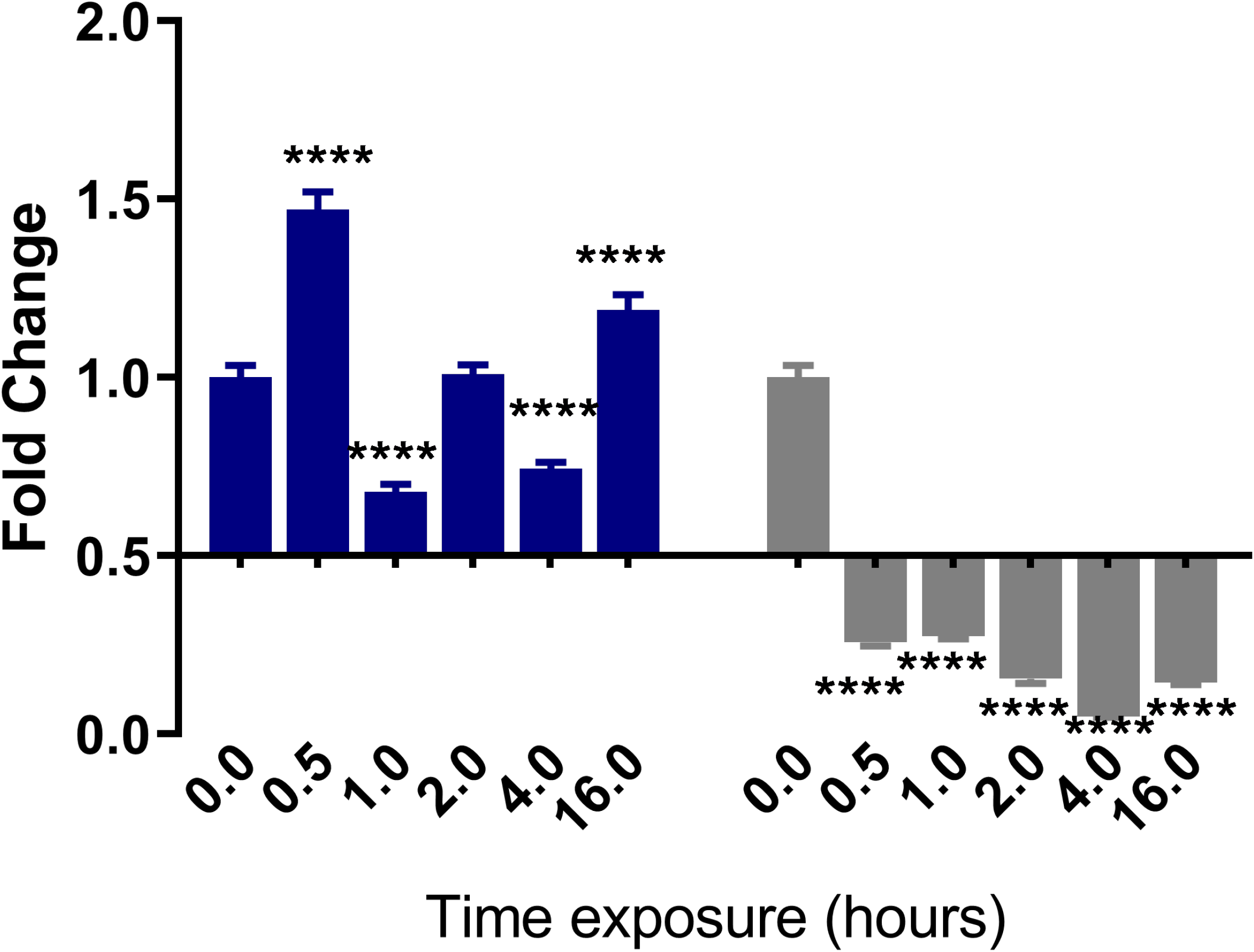
qPCR analysis of RNA from either resistant BS-90 (blue histogram) or susceptible BBO2 (gray histograms) juvenile snails unexposed (0) or exposed for increasing intervals (30 seconds to 16 hours) to *S. mansoni* miracidia. Histograms show expression of the *BgHAT* encoding transcript in snails at each time point from five biological replicates. Note the increase in fold change in the resistant BS-90 compared to the susceptible BBO2 snails after parasite infection. Significant expression normalized against expression of the myoglobin encoding transcript was measured by 2-way ANOVA and is indicated by number of asterixis on each histogram where ****, indicates the most significant value *p* ≤ 0.0001, *** *p* ≤ 0.001, ** *p* ≤ 0.01, * *p* ≤ 0.05, ns *p* > 0.05.

**Figure 1E.**
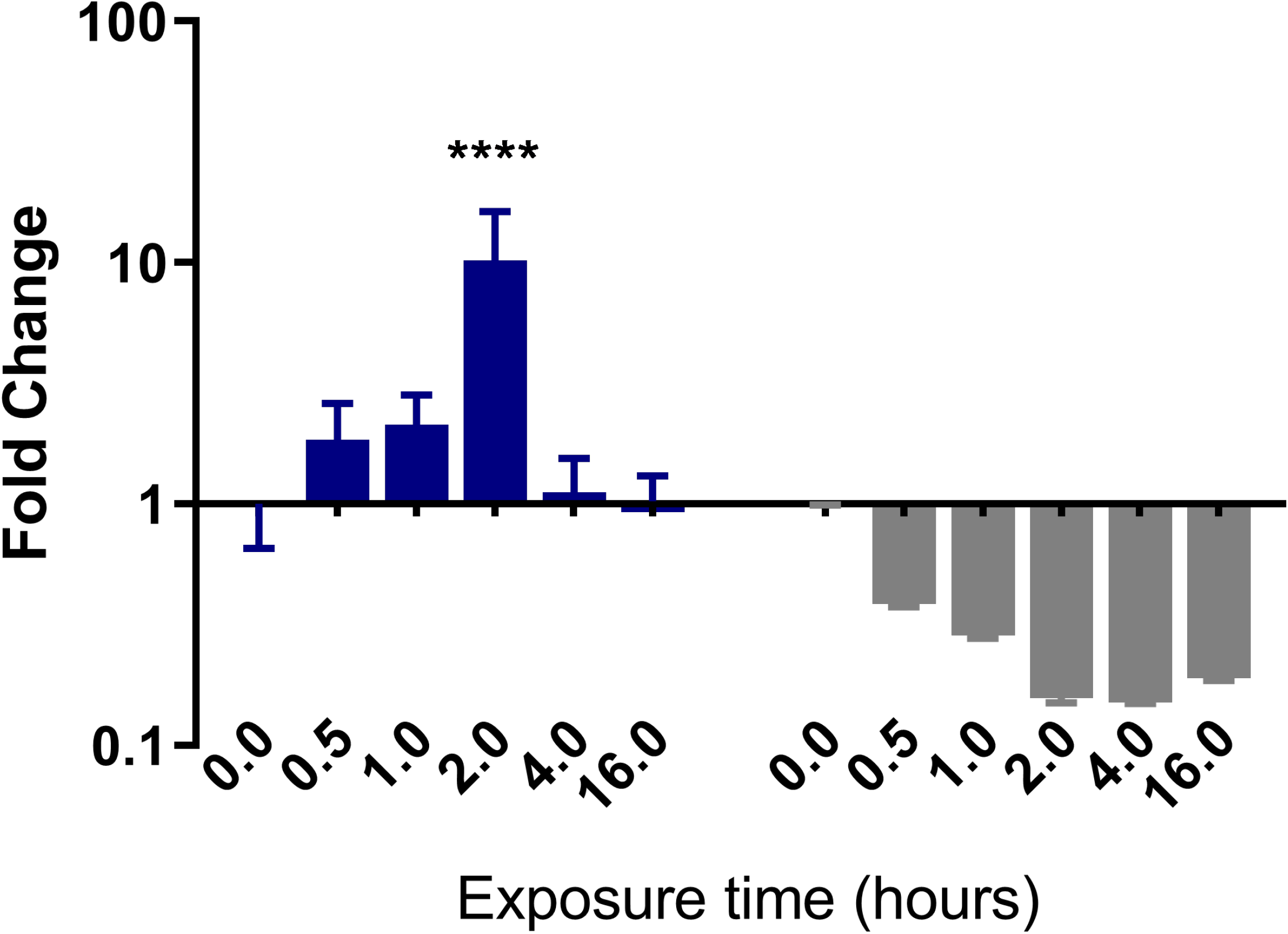
qPCR analysis of RNA from either resistant BS-90 (blue histogram) or susceptible BBO2 (gray histograms) juvenile snails unexposed (0) or exposed for increasing intervals (30 seconds to 16 hours) to *S. mansoni* miracidia. Histograms show expression of the *BgMT* encoding transcript in snails at each time point from five biological replicates. Note the increase in fold change in the resistant BS-90 compared to the susceptible BBO2 snails after parasite infection. Significant expression normalized against expression of the myoglobin encoding transcript was measured by 2-way ANOVA and is indicated by number of asterixis on each histogram where ****, indicates the most significant value *p* ≤ 0.0001, *** *p* ≤ 0.001, ** *p* ≤ 0.01, * *p* ≤ 0.05, ns *p* > 0.05.

### siRNA corresponding to *BgPiwi* knock down expression of the piwi encoding transcript rendering resistant BS90 snails susceptible

Since the findings revealed that the transcript encoding *BgPiwi* was upregulated in juvenile resistant BS90 snails after *S. mansoni* infection, we proceeded to examine the modulation of this (*BgPiwi*) transcript in the resistant (25°C) and the susceptible juvenile (32°C) BS90 snails, with and without *S. mansoni* infection. The findings presented in Figure 2A demonstrate that unlike resistant BS90 snails, cultured at 25°C, where expression of the *BgPiwi* encoding was upregulated following (2 hour) *S. mansoni* infection; in susceptible juvenile BS90 snails, cultured at 32°C, the *piwi* transcript was downregulated, similarly to *S. mansoni* infection of the susceptible BBO2 snail (Fig. 1A). Based on these data, and given he known role of PIWI in silencing endogenous retrotransposable elements and previous results that revealed expression (upregulation) of the RT domain of *nimbus* occurring in juvenile susceptible but not resistant snails in response to *S. mansoni*, the functional role of PIWI in the epigenetics of *B. glabrata*/*S. mansoni* susceptibility was investigated by silencing the expression of the transcript encoding *BgPiwi* by using corresponding siRNAs. As shown in Figure 2B, investigation of the expression of the transcript encoding *BgPiwi* in siRNA/PEI transfected snails with (2 hours) and without (0 hours) *S. mansoni* exposure showed the knock-down of the *BgPiwi* encoding transcript in snails transfected with siRNAs corresponding to *BgPiwi,* but not to the universal mock siRNA. Use of *Bgpiwi* dsRNA/PEI complexes instead of siRNA/PEI complexes to transfect BS90 snails, similarly, produced the knock-down of the *piwi* encoding transcript as observed with using siRNAs. To determine the biological effect of silencing *BgPiwi* in relation to *S. mansoni* infection in *BgPiwi* siRNA/PEI transfected BS90 snails, schistosome exposed transfected non-transfected normal snails and Universal mock siRNA/PEI transfected snails were left at room temperature and evaluated at 4-and 6-weeks post-exposure. As shown in Figures 2C and 2D, BS90, normally resistant snails, transfected with *BgPiwi* siRNA/PEI, shed cercariae at 4- and 6-weeks post-exposure to *S. mansoni*.

**Figure 2A.**
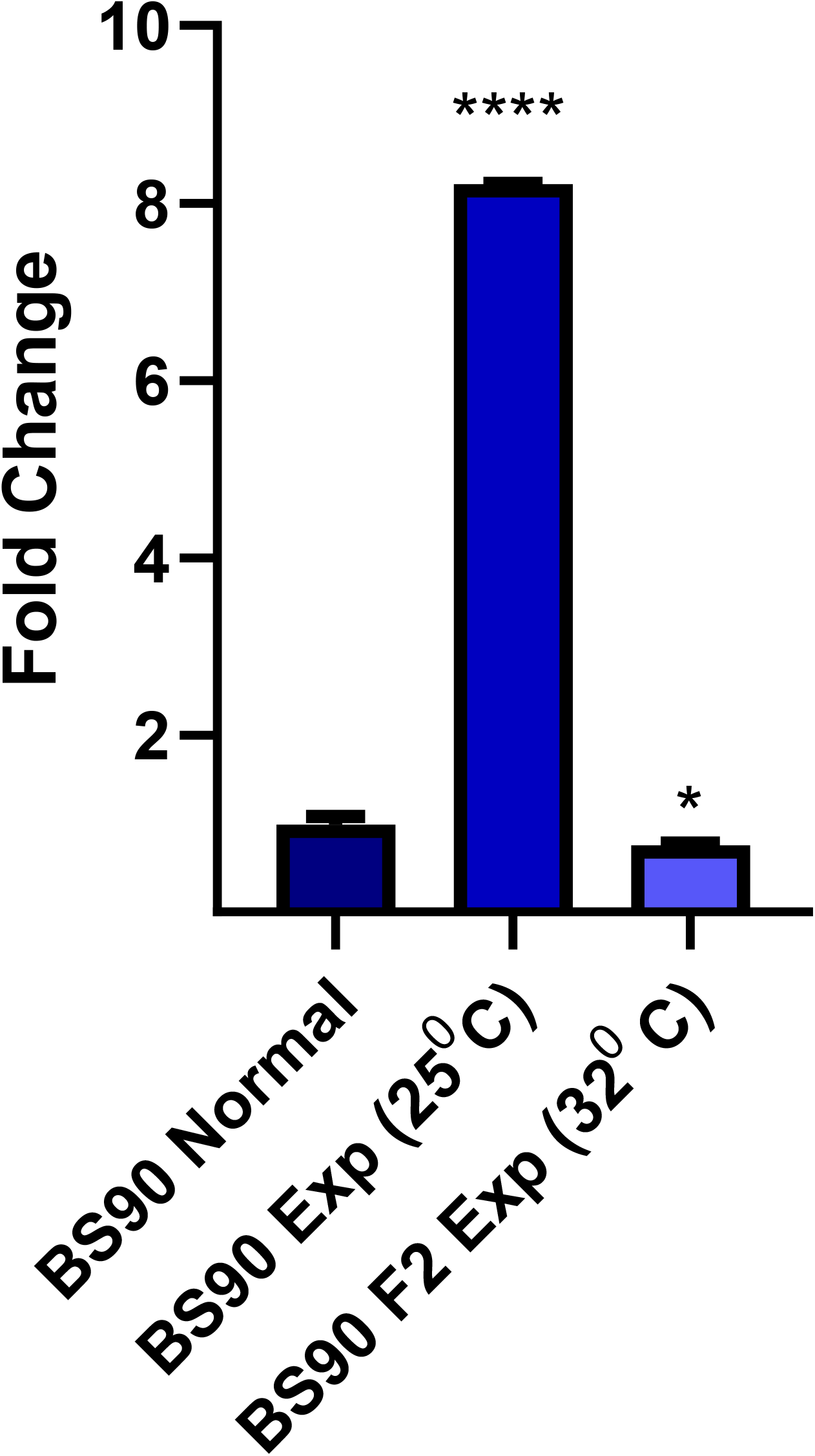
qPCR analysis of RNA from either non permissive (25°C) resistant BS-90 (blue histogram) or permissive (32°C) susceptible BS-90 (gray histograms) juvenile snails unexposed (normal) or exposed for 2hr to *S. mansoni* miracidia. Histograms show expression of the *BgPiwi* encoding transcript in these snails residing at different temperatures. Note the significant induction (8-fold change) in 25°C non-permissive BS-90 snails compared the down regulation of the transcript in permissive BS-90 snails residing at 32°C after parasite infection. Fold change was determined as described in materials and methods. Significant expression normalized against expression of the myoglobin encoding transcript was measured by 2-way ANOVA and is indicated by number of asterixis on each histogram where ****, indicates the most significant value *p* ≤ 0.0001, *** *p* ≤ 0.001, ** *p* ≤ 0.01, * *p* ≤ 0.05, ns *p* > 0.05.

**Figure 2B.**
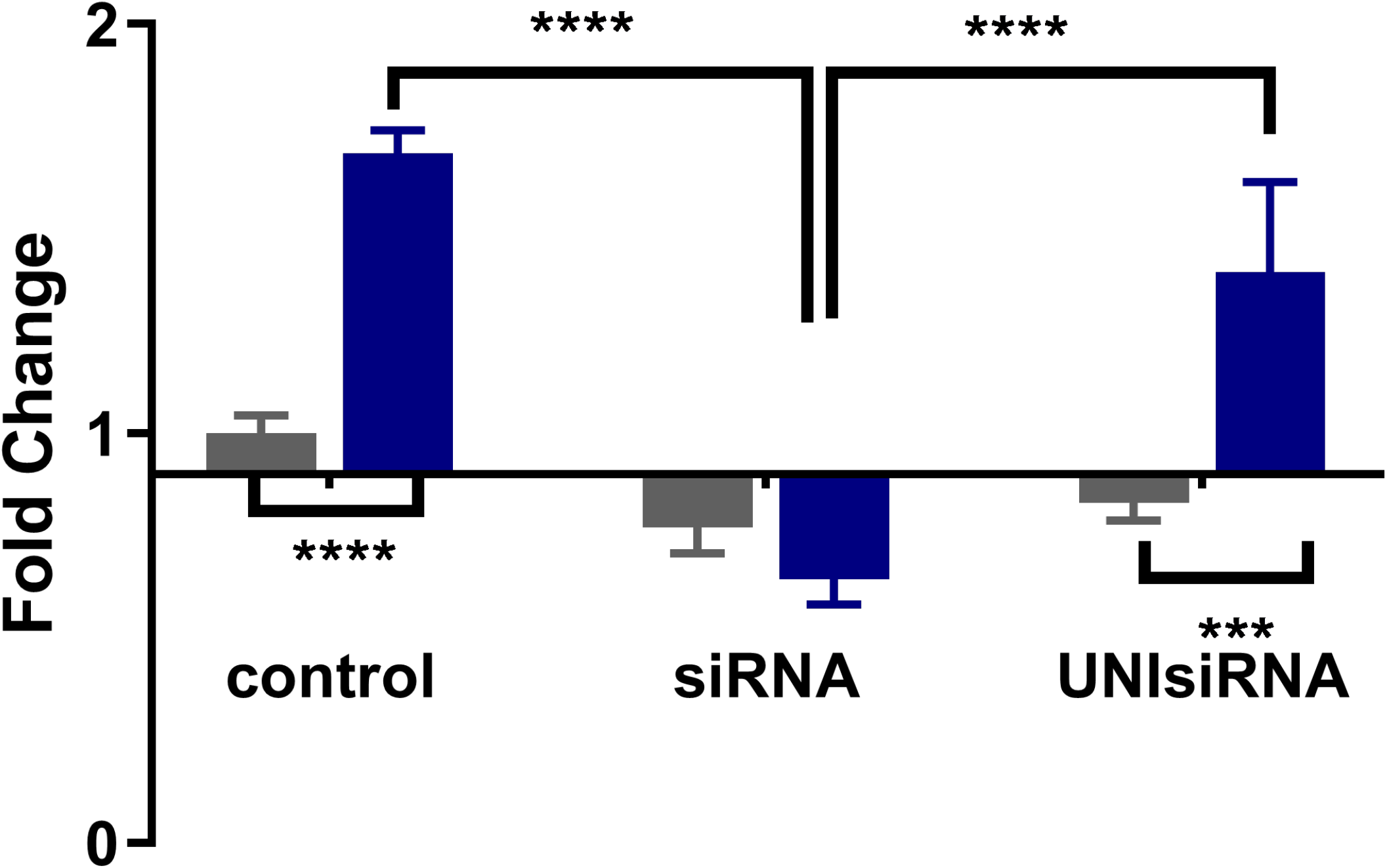
qPCR analysis of RNA from resistant BS-90 juvenile snails unexposed (gray) or exposed (blue) for 2hr to *S. mansoni* miracidia. Histograms show expression of the *BgPiwi* encoding transcript in normal BS-90 snails (control) or those transfected with *BgPiwi* siRNA. Note induction of the *BgPiwi* encoding transcript occurs in *S. mansoni* exposed control BS-90 snails and the knock down of the transcript in BS-90 (exposed and unexposed) snails transfected with *BgPiwi* siRNA. In BS-90 snails transfected with mock UNIsiRNA, note the upregulation of the *BgPiwi* encoding transcript in exposed snails similar to induction observed in control exposed snails. Fold change was determined as described in materials and methods. Significant expression normalized against expression of the myoglobin encoding transcript was measured by 2-way ANOVA and is indicated by number of asterixis on each histogram where ****, indicates the most significant value *p* ≤ 0.0001, *** *p* ≤ 0.001, ** *p* ≤ 0.01, * *p* ≤ 0.05, ns *p* > 0.05.

**Figure 2C.**
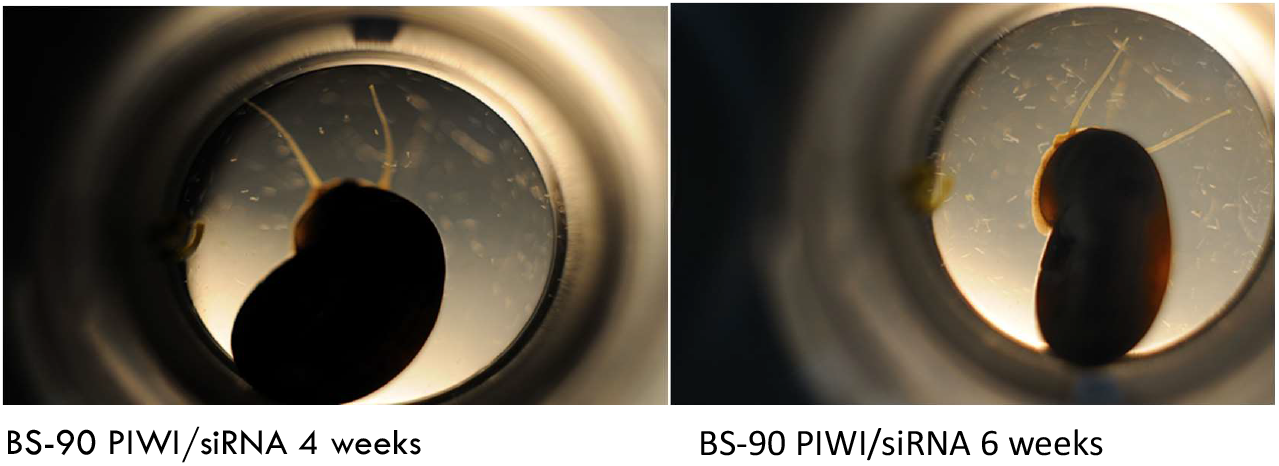
To determine the biological effect of silencing *BgPiwi* in relation to *S. mansoni* infection in *BgPiwi* siRNA/PEI transfected BS-90 snails, schistosome exposed *BgPiwi* siRNA transfected and non-transfected (not shown) snails were left at room temperature and evaluated at 4- and 6- weeks post-exposure. Note that the BS-90 snail transfected with *BgPiwi* siRNA shed cercariae at 4- and 6- weeks post-exposure to *S. mansoni*.

### Knock-down of piwi encoding transcript with *siBgpiwi*/PEI complexes concurrently upregulates the *nimbus* RT in transfected BS90 snails

Because PIWI suppresses the expression of retrotransposable elements, such as the *B. glabrata* non-LTR retrotransposable element *nimbus,* the transcription of the RT domain of this element was investigated in either unexposed control (0) or *S. mansoni* (2 hour) exposed BS90 snails that were transfected with either *Bgpiwi* siRNA or mock universal siRNA (Fig. 3A). The same cDNA templates utilized in the qPCR assays shown in Figure 2B were utilized in the analysis of the expression of *nimbus* RT (Fig. 3A). Using gene specific primers corresponding to the RT domain of *nimbus* results showed that in normal BS90 snails that were unexposed (0) to *S. mansoni,* that similar to resistant BS90 that were transfected with mock siRNA (UnisiRNA/PEI) expression of *nimbus* RT remained low upon infection of the normal resistant BS90 snails as demonstrated previously [18]. However, in BS90 snails transfected with *Bgpiwi* siRNA before exposure, the *nimbus* RT encoding transcript was upregulated when *Bgpiwi* transcript was silenced.

**Figure 3.**
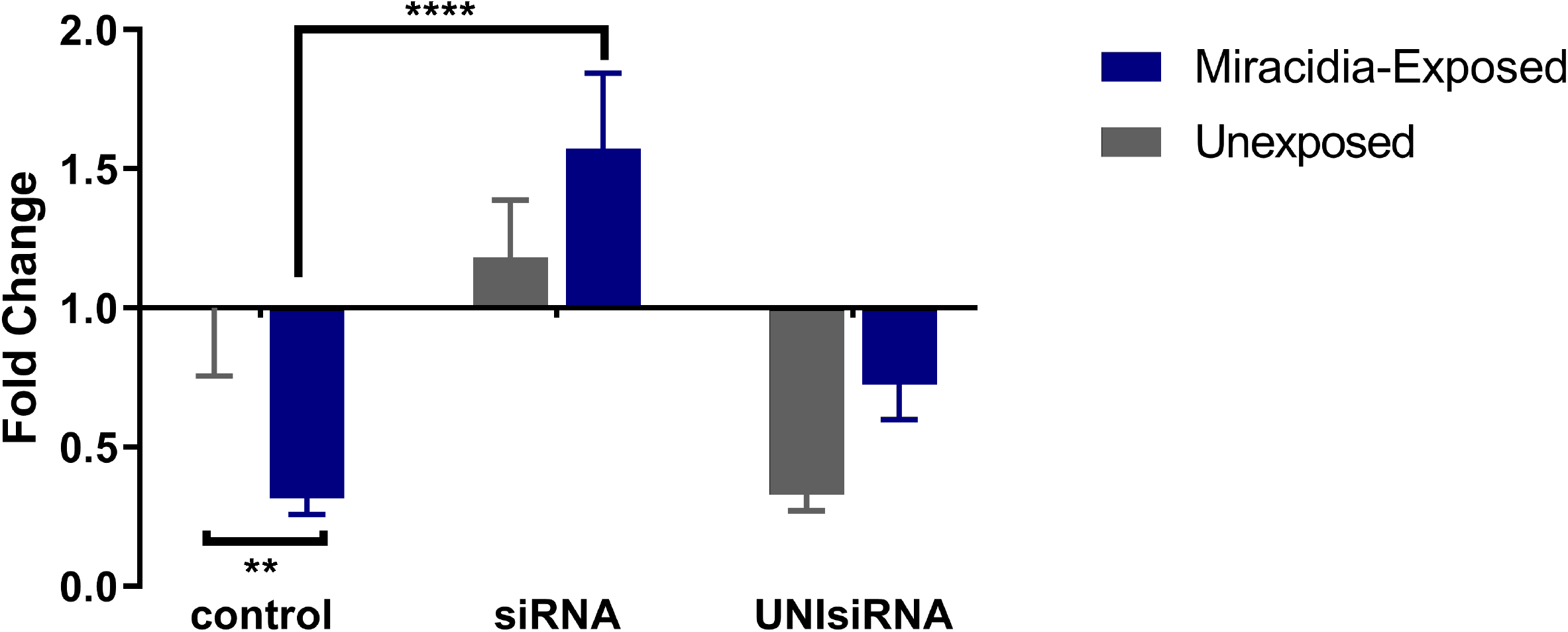
qPCR analysis of RNA from resistant BS-90 juvenile snails unexposed (gray) or exposed (blue) for 2hr to *S. mansoni* miracidia. Histograms show expression of the *nimbusRT* encoding transcript in normal BS-90 snails (control) or those transfected with *BgPiwi* siRNA. Note the down regulation of the *nimbusRT* encoding transcript in *S. mansoni* exposed control BS-90 snails and the upregulation of *nimbusRT* transcript in BS-90 (exposed and unexposed) snails transfected with *BgPiwi* siRNA where transcript encoding *BgPiwi* has been knocked-down (shown in Fig.2B). In BS-90 snails transfected with mock UNIsiRNA, note the down regulation of the *nimbusRT* encoding transcript in exposed snails similar to that observed in control exposed snails. Fold change was determined as described in materials and methods. Significant expression normalized against expression of the myoglobin encoding transcript was measured by 2-way ANOVA and is indicated by number of asterixis on each histogram where ****, indicates the most significant value *p* ≤ 0.0001, *** *p* ≤ 0.001, ** *p* ≤ 0.01, * *p* ≤ 0.05, ns *p* > 0.05.

### Lamivudine RT inhibitor treatment of susceptible BBO2 snails differentially regulates transcription of *nimbus* RT and *Bgpiwi*

To further examine the interplay between the expression of *Bgpiwi* and *nimbus* RT encoding transcripts in relation to *S. mansoni* infection of *B. glabrata*. The susceptible juvenile BBO2 snail was treated with the RT inhibitor drug, lamivudine, as described in materials and methods prior to exposure to *S. mansoni.* As shown in Figure 4, panels A and B, qPCR analysis of the same cDNA template prepared from BBO2 susceptible snails that were either treated or untreated with different concentrations (100 ng/ml and 200 ng/ml) of lamivudine before exposure (for 2 hours) or not exposed (0 hour) to *S. mansoni* were performed by utilizing primers corresponding to the *Bgpiwi* (4A) or *nimbus* RT (4B) encoding transcripts. Figure 4A shows the expression of *Bgpiwi* remained relatively unchanged with drug treatment while that of *nimbus* RT (Fig. 4B) was downregulated with Lamivudine. Both concentrations of lamivudine, either 100 or 200 ng/ml, used to treat BBO2 snails prior to exposure knocked-down the expression of the *nimbus* RT encoding transcript in lamivudine–treated snails.

**Figure 4A.**
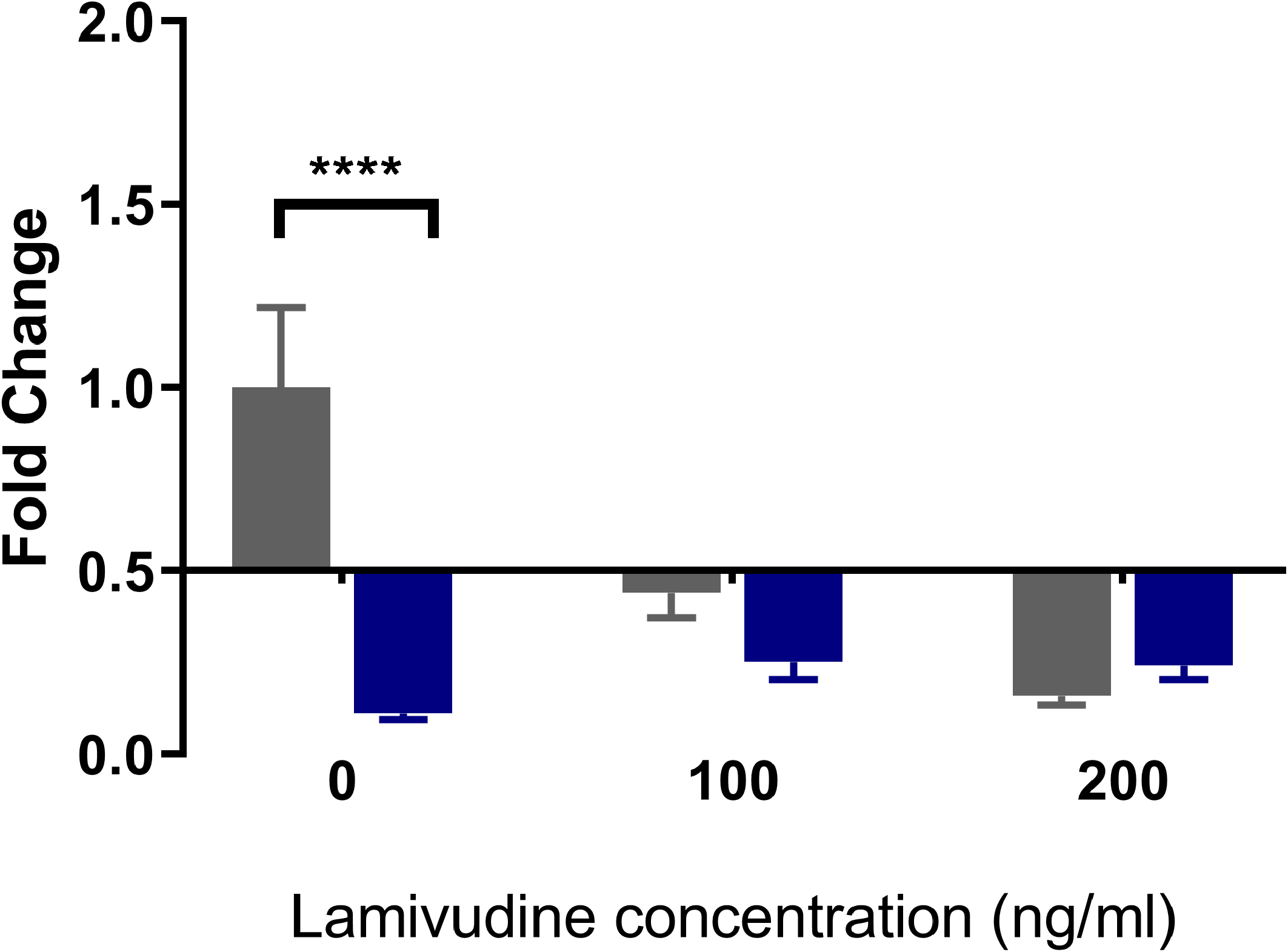
qPCR analysis of RNA from susceptible BBO2 juvenile snails unexposed (gray) or exposed (blue) for 2hr to *S. mansoni* miracidia. Histograms show expression of the *BgPiwi* encoding transcript in normal BBO2 snails (0) or those treated with RT inhibitor Lamivudine (100 ng/ml or 200ng/m1). Note the down regulation of the *BgPiwi* encoding transcript in exposed control (0) snails and lamivudine treated snails (exposed and unexposed) snails. Fold change was determined as described in materials and methods. Significant expression normalized against expression of the myoglobin encoding transcript was measured by 2-way ANOVA and is indicated by number of asterixis on each histogram where ****, indicates the most significant value *p* ≤ 0.0001, *** *p* ≤ 0.001, ** *p* ≤ 0.01, * *p* ≤ 0.05, ns *p* > 0.05.

**Figure 4B.**
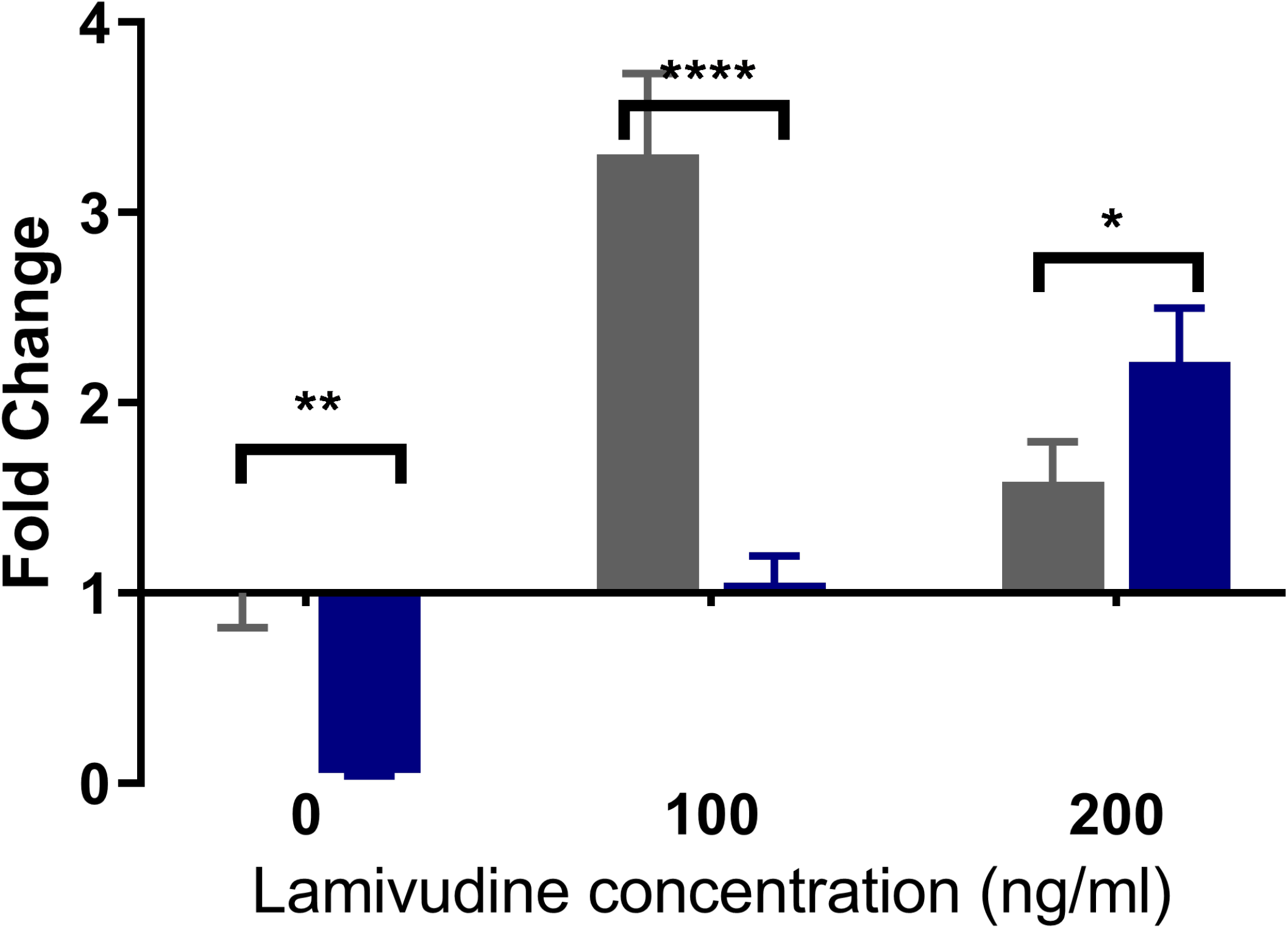
qPCR analysis of RNA from susceptible BBO2 juvenile snails unexposed (gray) or exposed (blue) for 2hr to *S. mansoni* miracidia. Histograms show expression of the *nimbusRT* encoding transcript in normal BBO2 snails (0) or those treated with RT inhibitor Lamivudine (100 ng/ml or 200ng/m1). Note the upregulation of the *nimbusRT* encoding transcript in exposed control (0) snails and down regulation of this transcript in Lamivudine treated snails (exposed and unexposed) snails. Fold change was determined as described in materials and methods. Significant expression normalized against expression of the myoglobin encoding transcript was measured by 2-way ANOVA and is indicated by number of asterixis on each histogram where ****, indicates the most significant value *p* ≤ 0.0001, *** *p* ≤ 0.001, ** *p* ≤ 0.01, * *p* ≤ 0.05, ns *p* > 0.05.

### Lamivudine blocks schistosome infection of BBO2 snails

Because the silencing of the *Bgpiwi* encoding transcript rendered the resistant BS90 snail susceptible, concurrent with the down- and up-regulation of the *Bgpiwi* encoding transcript and *nimbus* RT encoding transcripts, respectively, and since follow-up qPCR analysis using the same cDNA templates showed that *nimbus* RT was knocked down by lamivudine treatment of the susceptible BBO2 snail, the effect of this drug on the ability of the susceptible snail to sustain a viable infection was examined (Figure 5). As shown in Figure 5A, where juvenile BBO2 snails were either treated with 100 ng/ml of lamivudine either before or after two weeks exposure to *S. mansoni* miracidia, snails failed to shed cercariae when treated with lamivudine prior to exposure (Fig. 5A). However, snails treated with BPPA (anthraquinone diacetate), another RT inhibitor that specifically blocks the reverse transcriptase activity of the human homolog of telomerase (hTERT) in *B. glabrata* shed cercariae when treated before exposure but not when treated at 14 days post parasite exposure (Fig. 5B) in contrast to the outcome when snails were treated with lamivudine before *S. mansoni* infection. Treatment of the snails with lamivudine at day 14 post-exposure, however, showed some snails shed cercariae when the drug was utilized later after infection.

**Figure 5A.**
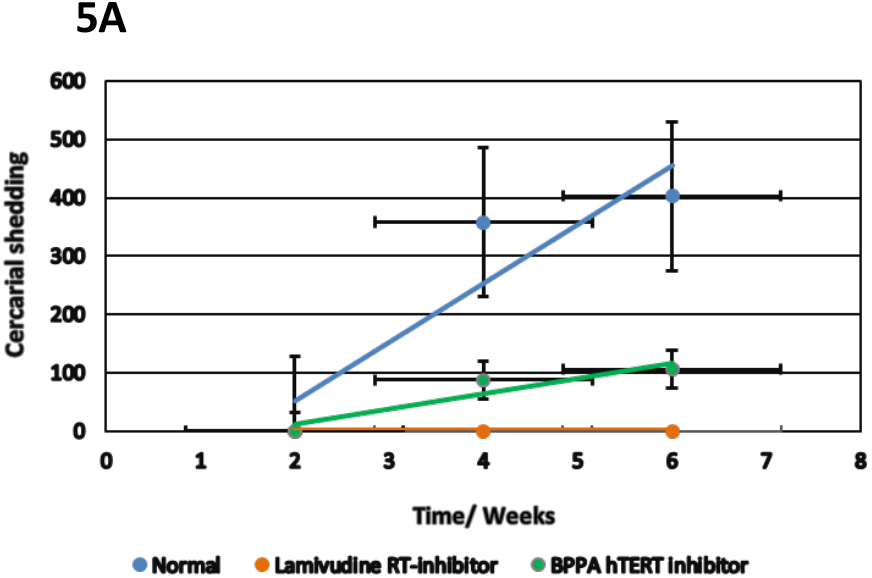
To determine the effect of lamivudine in relation to *S. mansoni* infection of susceptible BBO2 snails, snails were treated before exposure with 100 ng/ml of the RT inhibitor, maintained at room temperature, and evaluated for up to 6 weeks post-exposure. Note that the BBO2 snails treated before infection with lamivudine failed to shed cercariae at 6 weeks post-exposure to *S. mansoni* unlike in untreated (control) snails. Also note that BBO2 snails treated with the hTERT RT inhibitor BBPA before exposure, unlike Lamivudine shed cercariae at 6 weeks post-exposure.

**Figure 5B.**
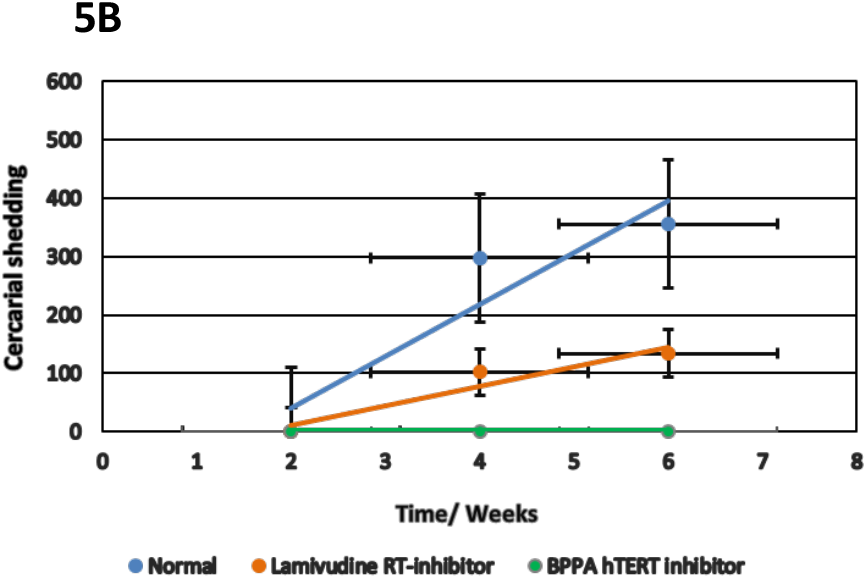
The effect of treating BBO2 susceptible snails with 100 ng/ml of lamivudine at 14 days post -exposure to *S. mansoni* was compared to the effect of treating susceptible BBO2 snails before exposure with 100 ng/ml of BPPA as described in materials and methods and left at room temperature and evaluated for up to 6 weeks post-exposure. Note that the BBO2 snails treated before *S. mansoni* exposure with 100 ng BPPA failed to shed cercariae at 6- weeks post-exposure unlike in untreated (control) snails. Also note that BBO2 snails treated with lamivudine at 14 days after infection shed cercariae at 6 weeks post-exposure.

### Relocalization of the *piwi* gene locus occurs in resistant *B. glabrata* snails upon infection

We had demonstrated that co-culture of snail Bge cells with live parasite [34, 36], and infection of whole snails with live parasite, both lead to non-random gene loci relocation within the interphase nuclei of the host snail cells, correlated with gene up-regulation [13, 19, 40], with notable differences in gene movement between susceptible and resistant snails. We have been able to demonstrate again that gene loci change their non-random nuclear location with changes in gene expression. A FISH probe containing the sequences for *B. glabrata piwi* was employed to delineate the nuclear position of the *piwi* gene loci in three snail strains, BS90 (resistant) and the two susceptible strains, BB02 and NMRI (Figure 6). The nuclear positioning of the gene loci were analyzed using the erosion analysis script for gene and chromosome positioning, we have used previously for different species[41–43] and can be assigned to a peripheral (Figure 6A), intermediate (Figure 6B) or internal (Figure 6C) location in cell nuclei. Notably, the *piwi* gene signal is in different nuclear compartments for the resistant and susceptible snails. Indeed, in BS90 (Figure 6D), the gene loci are found towards the nuclear interior and upon infection there is a relocation towards the nuclear periphery at 30 minutes after infection (Figure 6D), which coincides with the increase in *piwi* transcripts (Figure 1A). By two hours, the gene loci are relocating back to the nuclear interior (Figure 6D).

**Figure 6.**
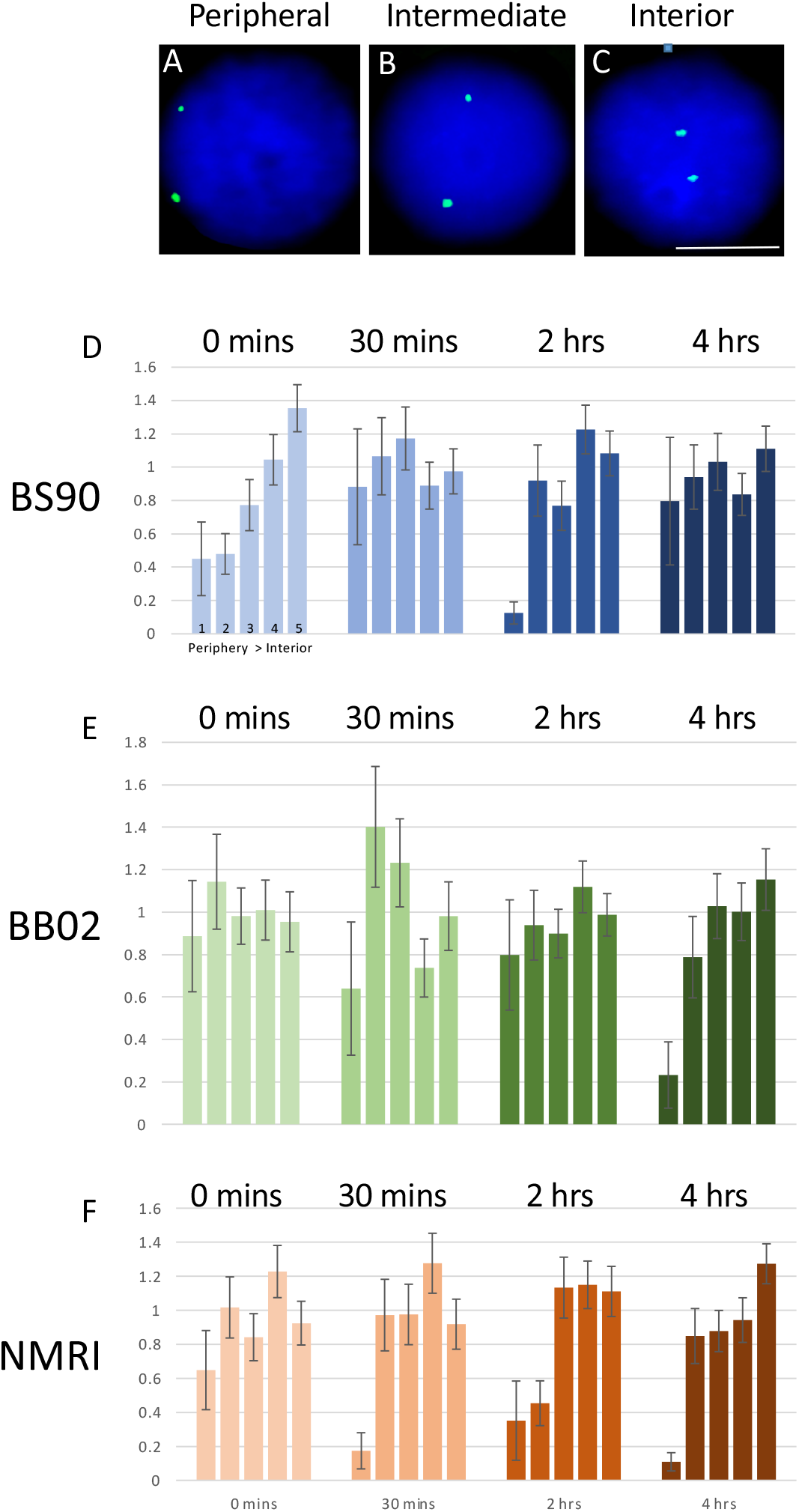
Nuclei (blue) were isolated from snail strains BS90, BB02 and NIMR ovo-testis and subjected to 2D-fluorescence in situ hybridisation (FISH) with labelled probes for the transposable element, *piwi* (green). Scale bar = 5 μm. Using a bespoke nuclear positioning script that creates five concentric shells of equal area, shells 1-5, with shell 1 being the nuclear periphery and shell 5 the nuclear centre, the percentage of fluorescent green gene signal is measured in each shell for over 50 nuclei and divided by the percentage of blue fluorescent signal for the DNA content (DAPI) in each shell. The data are averaged and plotted as bar charts with standard error of the mean (SEM).

The two susceptible snail strains of BB02 (Figure 6E) and NMRI (Figure 6F) display a similar gene loci location for *piwi*, an intermediate location, different from the resistant BS90 strain (Figure 7D). Furthermore, the *piwi* gene locus does not change location until 4 hours post-infection, where in both snail strains, the gene locus is in the nuclear interior, a location has been correlated with down-regulation of expression in BS90, and in BB02 (Figure 1A). Together, these findings further support the notion that the parasite is able to influence genome behavior within its host for its own advantage in an epigenetic mechanism, through functional spatial positioning.

## DISCUSSION

Significant progress has been made in recent years towards elucidating the molecular basis of *S. mansoni* resistance/susceptibility in the intermediate snail vector *B. glabrata.* From these studies, it has become clear that the snail and schistosome relationship is complex and highly variable [13, 20, 44]. This study was undertaken to determine the role of epigenetics in shaping the relationship between the snail and the schistosome. The genetics of this interaction is well known and several molecular determinants that underlie the innate defense anti-schistosome response in *B. glabrata* have been identified as have sequences that are linked in the snail genome to resistance [14, 21, 45]. Epigenetics describes the inheritance of a reversible phenotype that is not influenced by any change in the sequence of DNA. Transgenerational epigenetic inheritance induced by environmental changes have recently reported the role of PIWI and small piRNA in stress-induced genome modifications (see [46]). The impact of viral infection on the siRNA and Argonaute /PIWI pathway has mainly been reported in insects[47, 48]. However, little is known regarding genes and proteins involved in this insect Ago-2/RNAi antiviral/stress defense response [49].

Variation in *B. glabrata* susceptibility to *S. mansoni* was investigated by examining the regulation of key transcripts that play a role in epigenetics. Our approach was to use representative juvenile snail stocks that are either resistant (BS90) or susceptible (BBO2) to the NMRI strain of *S. mansoni*, to examine the temporal regulation of transcripts encoding Piwi (*BgPiwi*), chromobox protein homolog 1 (*BgCBx1*), histone acetyl transferase (HAT) histone deacetylase (HDAC) and metallotransferase (MT) were examined in these snail stocks within 30 min to 16hr post-exposure to *S. mansoni*. The differential expression of these transcripts was confirmed by comparing RNA-seq datasets that were generated from non-permissive juvenile BS90 snails, cultured at room temperature (25°C), and their permissive counterparts cultured for two generations at 32°C. The snail infections were carried out exclusively with juvenile snails. Thus, to further validate the differential expression of these transcripts, resistant juvenile BS90 and susceptible juvenile BBO2 snails were exposed to *S. mansoni* before using real-time qPCR to confirm their modulation pre- and post-parasite exposure. Transcripts encoding PIWI (*BgPiwi*), chromobox protein homolog 1 (*BgCBx1*), histone acetyl transferase (HAT) histone deacetylase (HDAC) and metallotransferase (MT) were upregulated (1.8 to 10-fold) in the resistant (BS90) snail as compared to their downregulation in the susceptible juvenile snail (BBO2). Upregulation of the majority of these transcripts occurred within the first 30 min of exposure to *S. mansoni*, peaking at 120 min before subsiding. In earlier reports, we showed that the regulation of the RT domain of the *B. glabrata* endogenous non-LTR-retrotransposable element, *nimbus,* was linked to the early differential stress response observed between juvenile resistant and susceptible snails [18]. Accordingly, induction of RT occurred concurrently with the upregulation of the transcript encoding Hsp70 in the susceptible but not the resistant snail to *S. mansoni*. Exposure of *B. glabrata* to irradiated attenuated miracidia, however, failed to induce these stress-related transcripts in early-infected juvenile susceptible snails [25]. Given those findings and those presented here showing that upregulation of the *BgPiwi* encoding transcript, occurs in resistant BS90 snails residing at room temperature (where they are resistant) but not in their susceptible counterparts residing at 32°C, we examined the expression of the *Bgpiwi* transcript more closely in relation to the early expression of *nimbus* in susceptible and resistant snails. To reiterate, changes in transcription regulation were evident within the 30 minutes of infection. In this regard, these novel findings differ from those described by others where molecular interactions between the snail and schistosomes were performed much later after miracidia penetration at which time the responses we now report might have waned.

The existence of several piRNA sequences in *B. glabrata* has been described but their role in a piwi-piRNA mediated anti-parasite defense mechanism in the snail remains elusive [50]. However, to determine whether a piwi gene-silencing mechanism that involves *nimbus* RT plays a role in blocking transmission of schistosomes in *B. glabrata*, we utilized a previously developed PEI-mediated soaking method to deliver two different *BgPiwi* corresponding duplex siRNAs, simultaneously, into the resistant BS90 snail, thereby knocking-down the expression of the PIWI encoding transcript. The RNAi suppression of PIWI transcription, rendered these *Bgpiwi* siRNA/PEI transfected resistant BS90 snails susceptible and thus able to shed cercariae. While the transcript encoding *Bgpiwi* was reduced in siRNA/PEI transfected snails, in contrast, the expression of *nimbus* RT-encoding transcript was upregulated, indicating that a gene silencing mechanism mediated by the interplay of piwi and modulation of the transcription of *nimbus* RT plays a major role in *B. glabrata* susceptibility to *S. mansoni*. To provide further support for these findings, the susceptible BBO2 snail was treated with lamivudine, a known RT inhibitor. Lamivudine is a RT nucleoside analog inhibitor that is used to treat hepatitis B and HIV/ AIDS [51]. Treatment of either susceptible BBO2 or NMRI snails prior to schistosome exposure, consistently, after several biological replicates blocked *S. mansoni* infection in the snail. By contrast, BPPA, another RT inhibitor that specifically targets the RT activity of telomerase, did not block infection in *B. glabrata* by *S. mansoni* when snails were treated before parasite exposure. However, in snails treated at 2 weeks after exposure, BPPA prevented infection unlike what was observed with this treatment regimen with lamivudine. These findings indicated that the mechanism of action of this *nimbus*/PIWI interplay occurs very early in the *S. mansoni* and *B. glabrata* interaction. In ongoing studies, we have shown that an hTERT homolog is absent in the *S. mansoni* genome with the snail ortholog showing significant identity at the amino acid level to the human enzyme.

Work is currently underway to determine if using the same siRNA mediated gene silencing strategy as described above will reveal the significance of the other transcripts identified in this study and their functional role in epigenetics of the *S. mansoni/B. glabrata* relationship. Previously, we showed that hypomethylation of the stress Hsp70 protein locus precedes the early upregulation of the Hsp70 encoding transcript in *S. mansoni* exposed susceptible (NMRI) but not resistant (BS90) juvenile snails [52]. The findings here that the transcript encoding MT was upregulated in juvenile BS90 resistant but not susceptible BBO2 upon early parasite infection snails supports this earlier result [52]. Ideal follow up experiments to further verify the involvement of all the transcripts identified in this study in epigenetics of snail plasticity to *S. mansoni* susceptibility will be to edit their CDS by a permanent gene mutation such as by CRISPR/Cas9 to knockout their function. These approaches can likely contribute to deciphering epigenetic processes that underlie the susceptibility of the snail to the parasite. Toward this objective, molecular toolkits for gene editing in *B. glabrata* are being developed. CRISPR gene editing has been used to edit the allograft inflammatory factor gene in the *Bge* embryonic cell line from *B. glabrata* [53] and is finding utility in editing the schistosome genome [54] [55]. Confirming earlier findings, we show that in intact juvenile resistant and susceptible snails, within a short period post-exposure to *S. mansoni*, the non-random movement of the *piwi* locus within interphase nuclei in relation to its active transcription depending on the susceptibility phenotype of the snail)[19]. We have shown for the schistosome mediated relocation of gene loci that movement in interphase nuclei (from peripheral to interior location) occurs early after infection of susceptible, not resistant snails, and precedes transcription of the gene loci in question. A soluble factor(s) within excretory secretory products (ESPs) from the wild type miracidium that mediates the systemic reorganization of the host genome is yet to be uncovered. Aside from schistosomes, viruses are the only other pathogens that have been shown to also mediate non-random gene relocation. However, schistosomes are the first metazoan parasites that have been shown to possess the ability to manipulate the genome of the host in this profound spatio-epigenetic fashion.

To conclude, although a molecular basis exists for *B. glabrata* susceptibility to *S. mansoni*, it remains far from clear what precise pathways/mechanisms are responsible for parasite survival or rejection in the early infected snail. This is the first study to show that epigenetics, involving the interplay of PIWI and the endogenous non LTR-retrotransposable *nimbus*, plays a role in the plasticity of snail susceptibility to *S. mansoni.*

## Supporting information

Accession numbers and primer sequences

## ACKNOWLEDGEMENTS

This work was supported in part by funds from the Clement B.T. Knight Cancer foundation and a grant from the National Science Foundation Grant (Award No. 1622811). Travel awards to master’s students (O.A, S.B, and N.P) from the UDC foundation is also acknowledged. We thank Ms. Oumsilama Elhelu for her technical help and Dr. April Massey for her help and support in allowing Dr. Michael Smith to conduct his research for his Ph.D. dissertation at UDC. DH was partially supported by funds from the College of Health, Medicine and Life Sciences at Brunel University London.

## Notes

https://www.ncbi.nlm.nih.gov/sra/PRJNA687288

### Competing Interest Statement

The authors have declared no competing interest.

