## Supplementary material for "PIWI silencing mechanism involving the retrotransposon *nimbus* orchestrates resistance to infection with *Schistosoma mansoni* in the snail vector, *Biomphalaria glabrata*": Accession numbers and primer sequences

Table 1

|  | **Description** | **Sequence** | **Accession Number** |
| --- | --- | --- | --- |
| *1.* | *BgPIWI* | 5’:TTGCAAAATGGGCGGTGAAG | XP_013081375 |
|  |  | 3': TGACGAACTGACTGGCTCAC |  |
| 2. | *BgHDAC* | 5':CCACATAAGGCCACAGCAGA | XP_013075422.1 |
|  |  | 3': TAGTACTTGCCCTTGCCTGC |  |
| 3. | *BgCBX1h-1* | 5': CAACGTGCATTTAAGGCGGA | XP_013064838.1 |
|  |  | 3': CACTGCTGTCTGTAGCACCT |  |
| 4. | *BgHAT* | 5': CGGCGGCATTTATCTTGGTG | XP_013062341.1 |
|  |  | 3': TGTCAATGTGGCGTCGAAGA |  |
| 5. | *BgMT* | 5': CATAGTCCGGTTGGTGCAGA | XP_013081375 |
|  |  | 3':GCAGTTGGTAGCAGCAAGAGA |  |
| 6. | *nimbusRT* | 5’:GCTCCATTAAACCGAACAGAC | EF413179 |
|  |  | 3':CCCCGTAGATCATTGCTAAC |  |

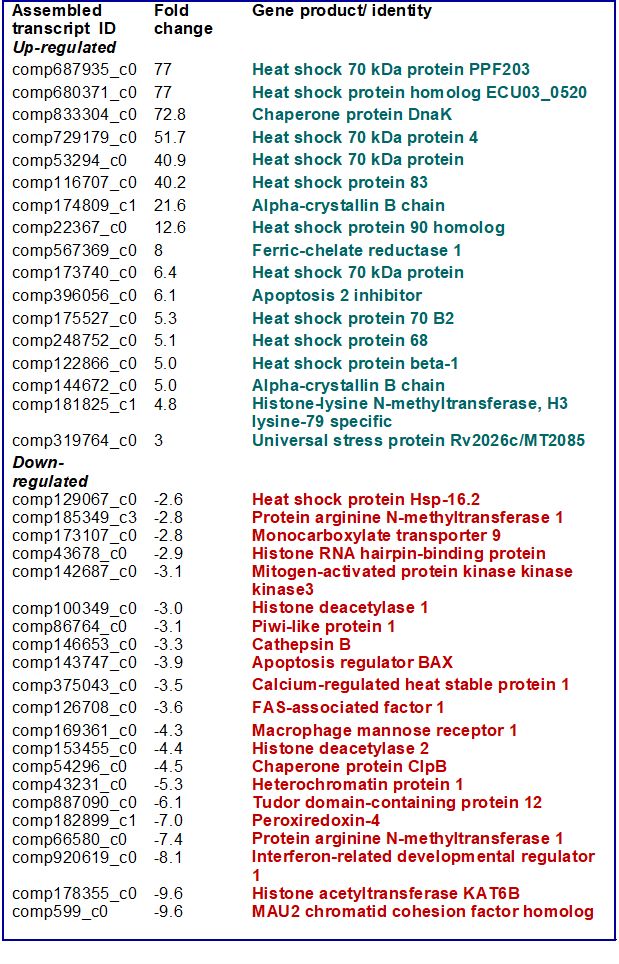
